# Tracking multiple genomic elements using correlative CRISPR imaging and sequential DNA FISH

**DOI:** 10.1101/101444

**Authors:** J. Guan, H. Liu, X. Shi, S. Feng, B. Huang

## Abstract

Live imaging of genome has offered important insights into the dynamics of the genome organization and gene expression. The demand to image simultaneously multiple genomic loci has prompted a flurry of exciting advances in multi-color CRISPR imaging, although color-based multiplexing is limited by the need for spectrally distinct fluorophores. Here we introduce an approach to achieve highly multiplexed live recording via correlative CRISPR imaging and sequential DNA fluorescence *in situ* hybridization (FISH). This approach first performs one-color live imaging of multiple genomic loci and then uses sequential rounds of DNA FISH to determine the loci identity. We have optimized the FISH protocol so that each round is complete in 1 min, demonstrating the identification of 7 genomic elements and the capability to sustain reversible staining and washing for up to 20 rounds. We have also developed a correlation-based algorithm to faithfully register live and FISH images. Our approach keeps the rest of the color palette open to image other cellular phenomena of interest, as demonstrated by our simultaneous live imaging of genomic loci together with a cell cycle reporter. Furthermore, the algorithm to register faithfully between live and fixed imaging is directly transferrable to other systems such as multiplex RNA imaging with RNA-FISH and multiplex protein imaging with antibody-staining.

## INTRODUCTION

Live imaging of genome has offered important insights into the dynamics of the genome organization and gene expression, both at the global nucleus scale (1, 2) and local chromatin scale (3, 4). Recent engineering efforts on DNA-binding protein systems have led to facile imaging of endogenous sequence-specific genomic loci in living cells (5, 6). The demand to image simultaneously multiple genomic loci has prompted a flurry of exciting advances in multi-color imaging methods where intriguing heterogeneous dynamics were observed for different loci. For example, in the CRISPR imaging systems, genomic loci are distinguished by labeling with different fluorescence proteins through Cas9 protein orthologues (7, 8) or modified single-guide RNA (sgRNA) scaffolds that recruit different RNA-binding proteins (9-12). In all these systems, the number of loci that can be distinguished simultaneously is still limited by the choice of fluorescence proteins that have sufficient color separation. Meanwhile, in fixed systems, highly multiplexed fluorescence *in situ* hybridization for both RNA (13-15) and DNA (16) has been reported by sequentially applying and imaging different probes following a prearranged code. Tens or even hundreds of DNA or RNA species can be distinguished in this way.

Here we report a correlative imaging method that combines the dynamic tracking capability of CRISPR imaging with the multiplicity of sequential FISH. This method allows us to perform live-cell CRISPR imaging first to obtain the dynamics of many genomic loci using one Cas9 protein and the corresponding sgRNAs followed by sequential rounds of DNA FISH to decode loci identity (Figure 1).

**Figure 1.**
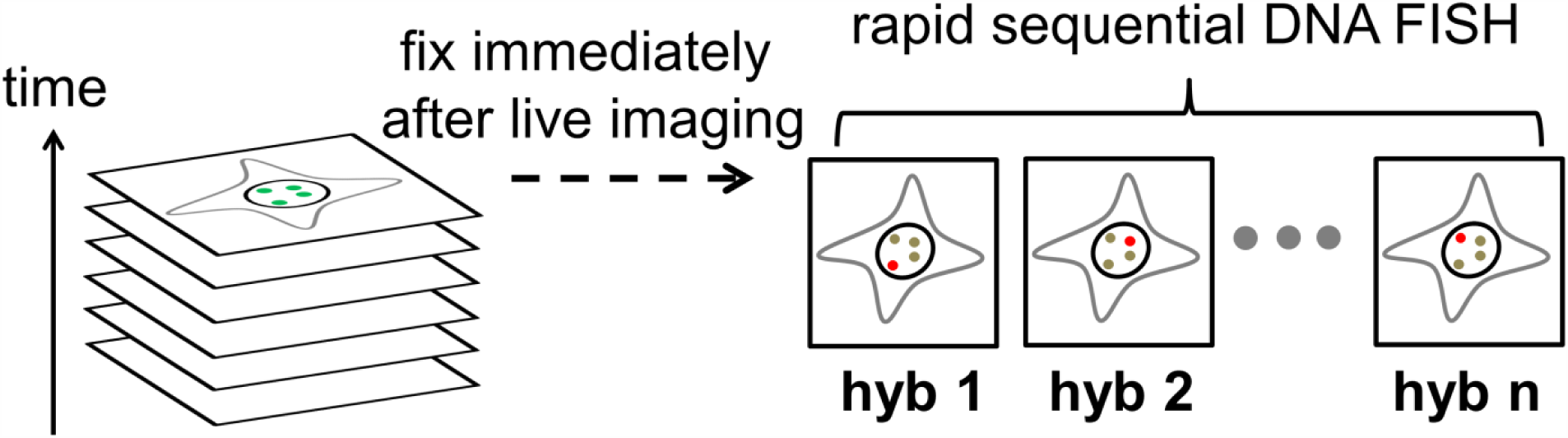
Schematic of correlative CRISPR imaging and sequential DNA FISH. Live cells are first imaged in time-lapse mode to acquire dynamics information. Multiple genomic loci are simultaneously imaged without distinguishing their identities. Cells are fixed immediately after live imaging. Rapid sequential rounds of DNA FISH are performed afterwards. As probes specifically bound to a locus are introduced in each round, the identity of the locus is resolved by comparing the last frame of live image and fixed images.

## MATERIALS AND METHODS

### Cell culture

Human retinal pigment epithelium (RPE) cells (ATCC, CRL-4000) were maintained in Dulbecco’s modified Eagle medium/Nutrient Mixture F-12 (DMEM/F-12) with GlutaMAX supplement (Gibco) in 10% Tet-system-approved fetal bovine serum (FBS) from Clontech. Human embryonic kidney (HEK) cell line HEK293T were maintained in DMEM with high glucose (UCSF Cell Culture Facility) in 10% Tet-system-approved FBS (Clontech). Cells were maintained at 37°C and 5% CO2 in a humidified incubator.

### Lentiviral production and stable expression of dCas9, sgRNA, and Fucci constructs

For viral production, HEK293T cells were seeded onto 6-well plate 1 day prior to transfection. 0.1 μg of pMD2.G plasmid, 0.8 μg of pCMV-dR8.91, and 1 μg of the lentiviral vector (Tet-on 3G, dCas9-EGFP, sgRNA, or Fucci) were cotransfected into HEK293T cells using FuGENE (Promega) following the manufacturer’s recommended protocol. Virus was harvested 48 hr post transfection. For viral transduction, cells were incubated with culture-medium-diluted viral supernatant for 12 hr. RPE cell lines stably expressing dCas9-EGFP were generated by coinfecting cells with a lentiviral cocktail containing viruses encoding both dCas9-EGFP and the Tet-on 3G transactivator protein (Clontech). Clonal cell lines expressing dCas9-EGFP were generated by picking a single-cell colony. The clones with low basal level expression of dCas9-EGFP were selected for CRISPR imaging. Clonal RPE cell line expressing dCas9-EGFP were transduced with lentivirus encoding Fucci (Gemini::RFP and Cdt1::mIFP linked by P2A domain) and cells with stable expression of Fucci was sorted using flow cytometry. The Fucci-containing cell line showed normal cell division and cell cycle progression.

### Optical setup and image acquisition

Fluorescence images were acquired on an inverted wide-field microscope (Nikon, Ti-E) with a 100× 1.45 N.A. oil immersion objective. The custom-build epi-illumination optics (Lumen Dynamics, X-Cite XLED1) provided excitation in DAPI, FITC, Cy3, and Cy5 channels. Quad-band dichroic excitation filter (Semrock, ZT405/488/561/640) was installed in the excitation path and quad-band emission filter (Semrock, FF410/504/582/669) in the emission path. Additional emission filters at 525 nm and 595 nm with 50 nm bandwidth were used for emission in FITC and Cy3 channels respectively to further reduce background noise. They were mounted into a motorized filter-wheel (Sutter Instrument, Lambda 10-B). A motorized microscope stage (ASI) controls the *xy* and *z* translation of the sample. The images were recorded with a sCMOS camera (Hamamatsu, C11440) in z-stacks of 6 μm with 0.3 μm steps. The microscope, light source, motorized stage, motorized filter wheel, and camera were controlled through custom configuration in Micro-Manager software. Live imaging was performed in FITC channel and FISH imaging was performed in Cy5 channel unless noted otherwise. Each image in Cy5 channel has a corresponding image in DAPI channel at the exactly same position to use in the two-step image registration algorithm described later. To improve mIFP signal, a final concentration of 25 μM biliverdin (Sigma 30891) was added to Fucci-containing cells at 12 hr before live imaging in FITC, Cy3, and Cy5 channels.

### DNA FISH

The bottom coverglass surface of an 8-well imaging chamber (Nunc Lab-Tek II, Thermo Fisher) was coated with 0.01% poly-L-lysine solution (Sigma) solution for 15 minutes and rinsed with PBS buffer three times. Cells were allowed to attach to the coverglass surface for overnight. Cells were fixed with 4% paraformaldehyde solution (Chem Cruz) at room temperature for 5 min followed by PBS buffer wash for three times. Cell membrane and nucleus membrane were permeabilized by methanol incubation for 5 min followed by PBS buffer wash. Cells were then heated on a hot plate at 80°C for 10 min in 80% formamide (Sigma). Cells were incubated for 2 min in hybridization solution of 200 nM oligo probes in the presence of 50% formamide and 2× SSC followed by PBS buffer wash three times. The conventional hybridization reagents such as dextran sulfate and blocking DNA reagents were not required. Imaging was performed in imaging buffer containing glucose, glucose oxidase, and catalase to prevent photobleaching. After each round of imaging in sequential FISH, the cells were washed with 80% formamide at 50°C for 30 s to remove the bound oligo probes followed by a new round of probe hybridization.

### Two-step image registration algorithm

To improve the reproducibility of sample positioning during repetitive mounting and unmounting steps, we designed a 3D-printed stage adaptor (Fig. S1 in the Supporting Material) that ensures tight fit of 8-well chambers on microscope stage. To precisely registered the acquired live and fixed images, we first performed a stage registration step (Fig. S2 in the Supporting Material) that allows the original region to be found after the sample is put back onto microscope stage. The position of the sample cell in live imaging at the last frame is termed as “original.” The position of the sample cell immediately after it is put back on stage is considered as “initial.” Image of nuclei at the initial position was convoluted to the image of nuclei in the original position to calculate full-image correlation. The images were down-sampled (a fraction of pixels, for example, 1/10, in both *x* and *y*) to speed up the registration algorithm for near real-time feedback in sample positioning. The images were then converted to binary format and convoluted to calculate correlation. The peak position in correlation corresponded to the original position in live imaging and the compensatory stage displacement to return to the original position was calculated based on the pixel size. The algorithm takes only a few seconds to process two 1024×1024 images. The motorized stage was then translated to the “adjusted” position based on the input of translation displacement. The spatial precision of this algorithm depends on the down-sampling and is usually found within 1 μm, consistent with the precision expectation at this step because the distance between every 10 pixels in the *x* or *y* direction is about 1 μm. Typically, the “initial” position is within the field of view of the “original” position. However, if necessary, a much larger imaging region, on millimeter scale, could be rapidly scanned and tiled using Micro-Manager and ImageJ stitching plugin to find the “original” position. The algorithm works well based on similar features between images. For example, when nuclei shape can be acquired in live imaging (e.g. diffusive nuclear EGFP signal in the current system), the correlation could be done between the last frame of live imaging and fixed DAPI signal. Alternatively, on stage DAPI staining is performed before the sample is taken off stage. The algorithm works well for both a single slice and a *z*-stack of nuclei.

Second, a further refinement in image registration was applied in image analysis to register between CRISPR loci at the last frame of live-cell imaging and FISH spots (Fig. S3 in the Supporting Material). Based on the nuclei shape, images taken in live EGFP channels at the last frame and fixed DAPI channels were registered and then the same image registration operation was applied to overlay the last frame of live images and FISH images. The registration algorithm is a modified version of image registration (17) that accounts for sample rotation and achieves subpixel precision through up-sampled discrete Fourier transforms (DFT) cross correlation. Briefly, the algorithm rotates an image in 1-degree steps and estimates the two-dimensional translational shift to register with a reference image through calculating the cross-correlation peak by fast Fourier transforming (FFT). The highest peak corresponds to not only the two-dimensional translational shift estimate but also the optimal angle of rotation. A refined translational registration with subpixel precision is then achieved through up-sampling the DFT in the small neighborhood of that earlier estimate by means of a matrix-multiply DFT. Our algorithm registered the images to a precision within 1/10 of a pixel. As the algorithm operates on Fourier Transform in frequency domain, it remains robust in image registration when features are more densely distributed (Fig. S4 in the Supporting Material).

### Single particle tracking

The positions of the spots in cell nuclei in live images were determined in CellProfiler. The position information at different time points is linked to generate trajectories using custom-written MATLAB (The MathWorks, Natick, MA) codes.

### Measurement of loci intensity

Z-stack images were first projected to generate an image using maximum-z projection and intensity measurement is performed on the projected images using custom-written MATLAB codes. The peak intensity of the genomic loci puncta was measured as the peak value in the selected region of interest subtracting nuclear background. The nuclear background was calculated as the mean value in nucleus regions lacking detectable puncta.

### Target genomic loci

We use hg19 version of human genome. The regions involved in this study are Chr1: 2581275-2634211; Chr3: 195505721-195515533 (denoted as Chr 3); Chr3: 195199025-195233876 (denoted as Chr3*); Chr7: 158122661-158135328; Chr13: 112930813-112973591 Chr19: 44720001-44760001 (denoted as NR1); Chr19: 29120001-29160001 (denoted as NR2); ChrX: 30806671-30824818.

#### sgRNA protospacer sequence to image human genomic loci

sgChr3, 5’ GUGGCGUGACCUGUGGAUGCUG 3’

sgChr7, 5’ GCUCUUAUGGUGAGAGUGU 3’

sgChr13, 5’ GAAGGAAUGGUCCAUGCUUACC 3’

sgChrX, 5’ GGCAAGGCAAGGCAAGGCACA 3’

#### DNA FISH probe sequence to image human genomic loci

Chr1, 5’ CCAGGTGAGCATCTGACAGCC 3’

Chr3, 5’ CTTCCTGTCACCGAC 3’

Chr3*, 5’ CCACTGTGATATCATACAGAGG 3’

Chr7, 5’ CCCACACTCTCACCATAAGAGC 3’

Chr13, 5’ GGTAAGCATGGACCATTCCTTC 3’

ChrX, 5’ TTGCCTTGTGCCTTGCCTTGC 3’

Telomere, 5’ CCCTAACCCTAACCCTAA 3’

Centromere, 5’ ATTCGTTGGAAACGGGA 3’

#### Non-repetitive probe design and synthesis

Oligo pool library was designed such that seven modules were concatenated. Two sets of index primer pairs were used to amplify the entire oligo pool library or selectively a sub-library of oligos. A variable region was designed to cover a genomic region of interest. Typically, 200 probes tiling over 40-kb genomic region lead to detectable FISH signal. The 30-nucleotide variable region was flanked on one side by T7 promoter used in in-vitro transcription and on the other side by a reverse transcription primer sequence shared in the entire library. The sequence in the variable region was first designed in OligoArray 2.1 software using parameter set -n 20 -l 30 -L 30 -D 1000 -t 70 -T 90 -s 76 -x 72 -p 35 -P 80 -m “GGGG;CCCC;TTTTT;AAAAA”. Sequences with homology of 17 nucleotides or more to the human genome were detected with blast+ and removed. Sequence with homology of 14 nucleotides or more due to concatenation between variable region and reverse transcription primer were also removed. The index primers and reverse transcription primers were designed by first truncating sequences to 20-mer oligo library from a 25-mer random oligo library (18). The oligos with a melting temperature between 75°C and 83°C were selected. Sequences with homology of 11 nucleotides or more or 5 nucleotides or more homology to the 3’ end within the 20-mer oligo library were removed. Oligos without a G or C base in the last two nucleotides on the 3’ end were removed. Sequences with homology to T7 promoter sequence were removed. Non-repetitive probe sequence files for loci NR1 and NR2 are included in the Supporting Material.

The oligo pool library is synthesized by CustomArray Inc and amplified by limited cycle PCR. *In-vitro* transcription (NEB, E2040S) was performed and dsDNA was converted to RNA with an effective amplification of 200+ fold. The RNA was then converted back to ssDNA in reverse transcription reaction (ThermoFisher, EP0751).

## RESULTS

### Fast FISH staining of genomic DNA

In order to practically and efficiently perform multiple rounds of DNA FISH on the same sample, we sought to address the technical challenge that DNA FISH often requires many hours or overnight for probe hybridization, which is much longer compared to that needed for RNA FISH because of the double-stranded nature of DNA. As recent reports suggest oligo DNA probes can rapidly label RNA or single-stranded DNA motif on the timescale of 5-30 minutes (15, 16, 19-21), we speculate that a rapid binding of oligo probes directly to genomic DNA is possible after sufficient DNA denaturation to separate duplex strands. To simplify the initial test, we started with oligo DNA probes targeting the tandemly repetitive sequence (TRS) in the human genome (Fig. S5 in the Supporting Material) so only one FISH probe is needed (22). Figure 2a-d shows the kinetics of FISH staining targeting a TRS region. Under optimized conditions, staining is essentially complete in 1 min and acceptable signal-to-noise ratio is even achieved in as short as 0.5 min, in drastic contrast to the common practice of overnight incubation. The representative images of cells stained for different durations show that the nuclear background is consistently low throughout staining. The efficiency of FISH staining, calculated as the ratio of observed to expected number of FISH puncta, reaches almost 100% by 1 min (Fig 2d; Fig. S6 in the Supporting Material). The signal-to-noise ratio reaches ~30 fold for the ~800 copies and we thus estimate a lower detection limit to be ~40 probes (Fig. S7 in the Supporting Material), a number consistent with those reported in recent RNA FISH studies. The signal-to-noise, nuclear background, and FISH efficiency remain constant after 1 min so a wide time window of hybridization works well. To test whether the optimized FISH protocol could also be widely applied to label non-repetitive genomic sequences, we designed oligo DNA probe pools that tile non-repetitive genomic regions (23) and observed similarly efficient staining of these regions (Figure 2i-j). Typically, we observe two puncta in each cell nucleus as expected for the diploid cells. Thus, this method can rapidly detect aneuploidy in interphase cells and potentially report copy number variations with probes targeting specific genes of interest.

**Figure 2.**
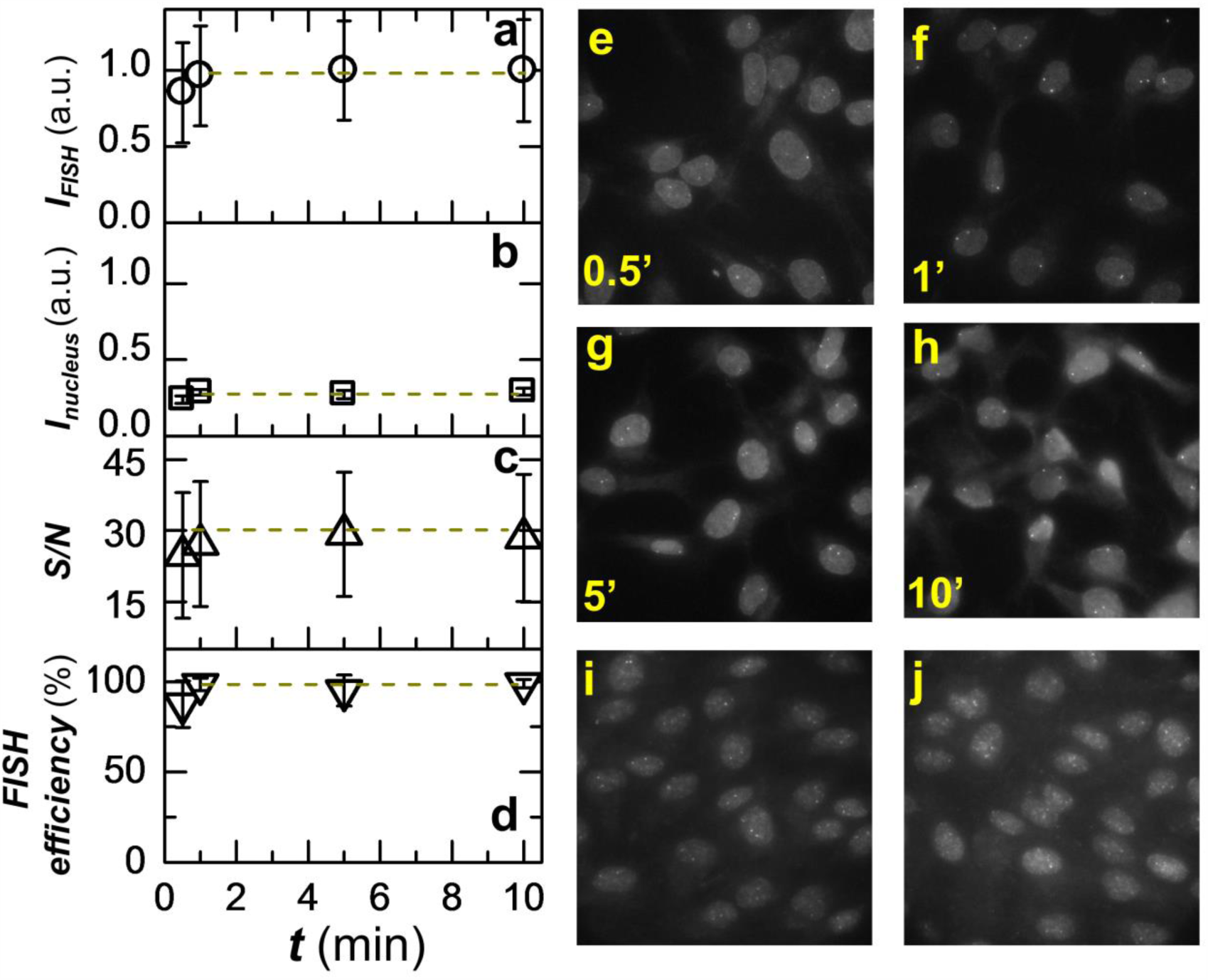
Rapid staining can be achieved with oligo DNA probes under optimized DNA FISH protocol. (a) Kinetics of the rapid DNA FISH staining. Intensity of FISH puncta for a tandemly repetitive sequence in the human genome is shown. Intensity is averaged over 300+ puncta and error bars denote standard deviation. (b) Intensity of nuclear background staining at different time points. (c) The signal-to-noise ratio of FISH puncta. (d) The efficiency of FISH staining, calculated as the ratio of observed to expected number of puncta. (e-h) Representative images at different time points of staining. Maximum intensity projection is shown for each 3D z-stack. (i-j) Representative images of two genomic regions denoted as NR1(b) and NR2 (c) of non-repetitive sequence labeled by tiling ~200 probes over 40-kb region with a staining time of 2 min. See Materials and Methods section for details on probe sequence. Maximum intensity projection is shown for each 3D z-stack.

Three aspects could have contributed to the fast staining: 1) gentle fixation by crosslinking and permeabilization by an alcohol wash dissolves most lipids in cell membrane and nuclear membrane; 2) an extended heating step ensures thorough denaturation between double strands of genomic DNA – we find that the genome accessibility directly correlates to the heating denaturation (Fig. S8 in the Supporting Material); and 3) oligo DNA allows extremely fast diffusion through cellular and nuclear structures. Note that the current FISH protocol is also greatly simplified in procedure, requiring no crowding reagents such as dextran sulfate to boost probe diffusion or blocking DNA reagents such as salmon sperm DNA and Cot-1 DNA to prevent non-specific nuclear staining.

### Multiple sequential rounds of DNA FISH in multiplex imaging

With the ease of performing rapid FISH staining, we then tested whether this technique can be applied to multiple sequential rounds of hybridization. A technical challenge here is to minimize the interference of bound probes with the next round of imaging. In previous studies, this issue was addressed by either DNase enzymatic reactions to degrade DNA probes bound to RNA targets (13) or photobleaching the dyes on bound probes using powerful lasers and then adding new probes targeting other vacant binding sites (15). Here to explore a simpler protocol, we apply a stringent wash step using concentrated formamide solution at elevated temperature to strip the bound oligo DNA probes after each round of imaging and detection. Figure 3 a-g shows an example of sequential FISH rounds where five distinct genomic regions, telomere, and centromere are sequentially detected through six rounds of staining and washing. The images show effective staining of specific sequences without interfering loci signals between sequential rounds. We find that the wash step is efficient and probes are effectively removed within 0.5 min. In Figure 3, we have also demonstrated two color FISH (Cy3 and Cy5 channel) to distinguish two loci as close as 300 kb away on chromosome 3 in one FISH round, suggesting the multiplex imaging capacity of the method could be further expanded through combining multi-color approaches.

**Figure 3.**
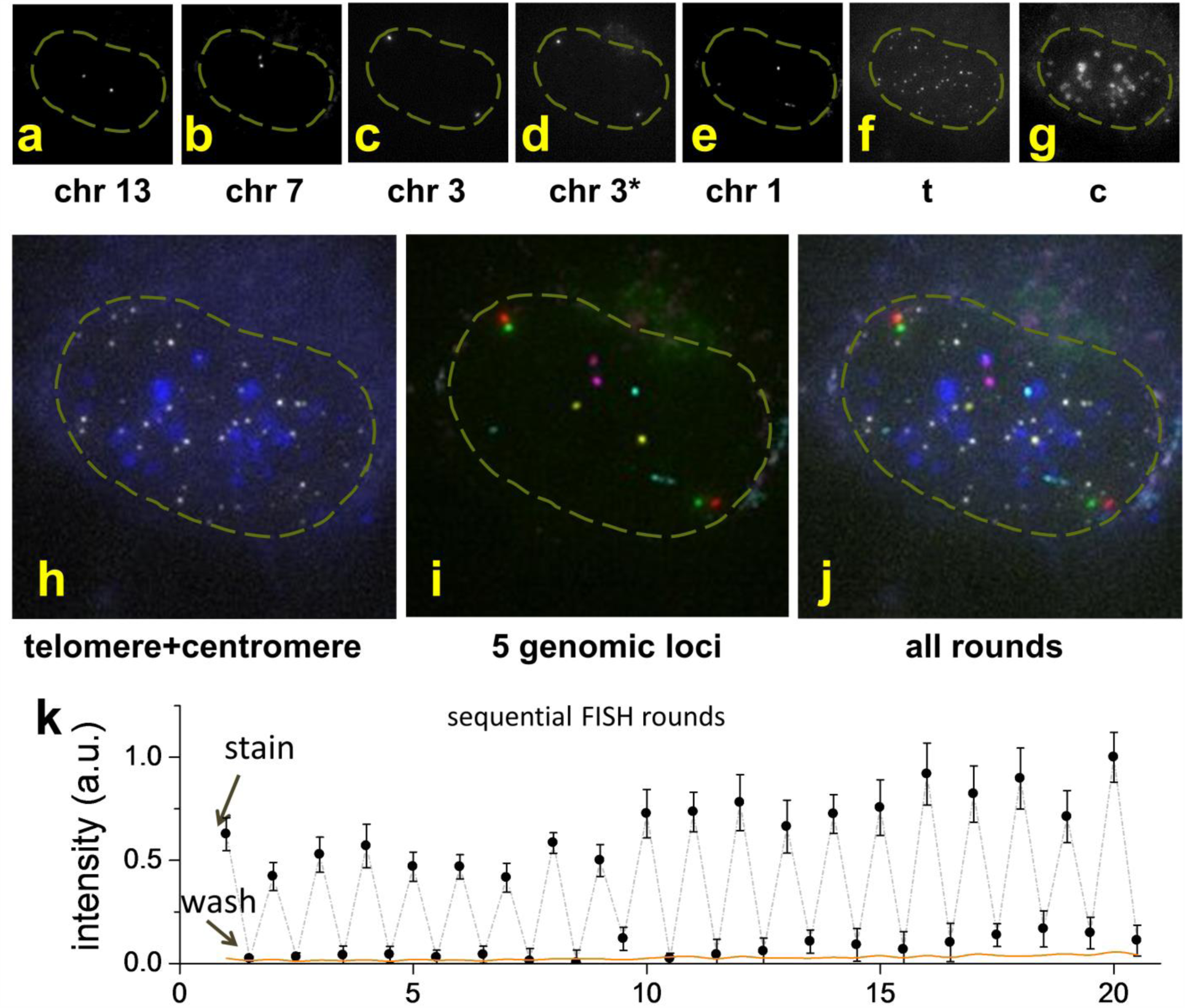
Multiple genomic loci can be resolved through multiple sequential rounds of DNA FISH. (a-g) Chr13, Chr7, Chr3, Chr3*, Chr1, telomere and centromere are sequentially resolved in six rounds of staining of 1 min and wash of 0.5 min. Two sites 300 kb apart on chromosome 3 (denoted as Chr3 and Chr3*) are co-stained and distinguished in one round with two colors. Maximum intensity projection is shown for each 3D z-stack. The nucleus contour is denoted as dashed yellow line. (h) False color overlay of telomere (white) and centromere (blue) staining. (i) False color overlay of Chr13 (yellow), Chr7 (magenta), Chr3 (red), Chr3* (green), and Chr1 (cyan). (j) Overlay of all loci determined in the sequential DNA FISH. Color is the same as in h and i. Image area in a-j is 22 μm×22 μm. (k) Peak intensity value of 20 rounds of repeated staining of 1 min and washing of 0.5 min with nuclear background subtracted, averaged over 8 loci. Error bars denote standard deviation. The nuclear background (orange line) is calculated as the average intensity within a nuclear region minus the average intensity from the empty region without cells.

Recent studies show that multiple rounds can quickly expand the multiplex capacity to thousands or more through various encoding strategies (13, 15, 16). To further test the potential of the method for multiplex imaging, we measured intensities of a specific genomic region through 20 rounds of alternating staining and washing. Figure 3k shows that the loci can be reversibly and consistently stained for at least 20 rounds without visible sample deterioration. High contrast is reproducibly seen with consistently high intensity after staining and almost negligible signal after wash. In fact, the residual signal after washing is so dim that it is often undetectable from nuclear background, especially in earlier rounds. The consistent and reversible staining and washing in multiple DNA FISH rounds suggest the multiplex potential of the method.

### Sequential DNA FISH after live imaging resolves loci identity

Figure 4a-d shows representative time-lapse images of a cell with simultaneous labeling of multiple genomic loci. Here, four sgRNAs with different protospacer sequence are expressed in the cell nucleus at the presence of dCas9::EGFP fusion protein, resulting in efficient labeling of the corresponding genomic loci and eight bright spots in the nucleus as homologous chromosomes are simultaneously labeled in the diploid cell. As these sgRNAs share the same dCas9 binding motif, one cannot directly distinguish the loci identities based on the live images. Nonetheless, genome dynamics can be extracted regardless of the loci identities. Comparison of images at different time points (Figure 4a-c) suggests that the relative positions of these loci are essentially stable, consistent with earlier FRAP measurement on cell nuclei (1, 2). The trajectory over 25 min overlaid on live images reveals a global motion of the nucleus (Figure 4d). The adjusted chromosome dynamics after subtracting the contribution of global nucleus movement show a more randomized trajectory direction (Fig. S9 in the Supporting Material). Furthermore, we show in Figure S10 that the global movement of nucleus is quite commonly seen, more so on longer observation timescale. We find that this global scale of nucleus movement is not caused by stage drift as cells in the same field of view have randomized trajectory direction with respect to one another. The live images also capture the dynamic vibrations between sister chromatids (arrowheads in Figure 4a), which are sometimes distinguishable when the distance exceeds the optical diffraction limit.

**Figure 4.**
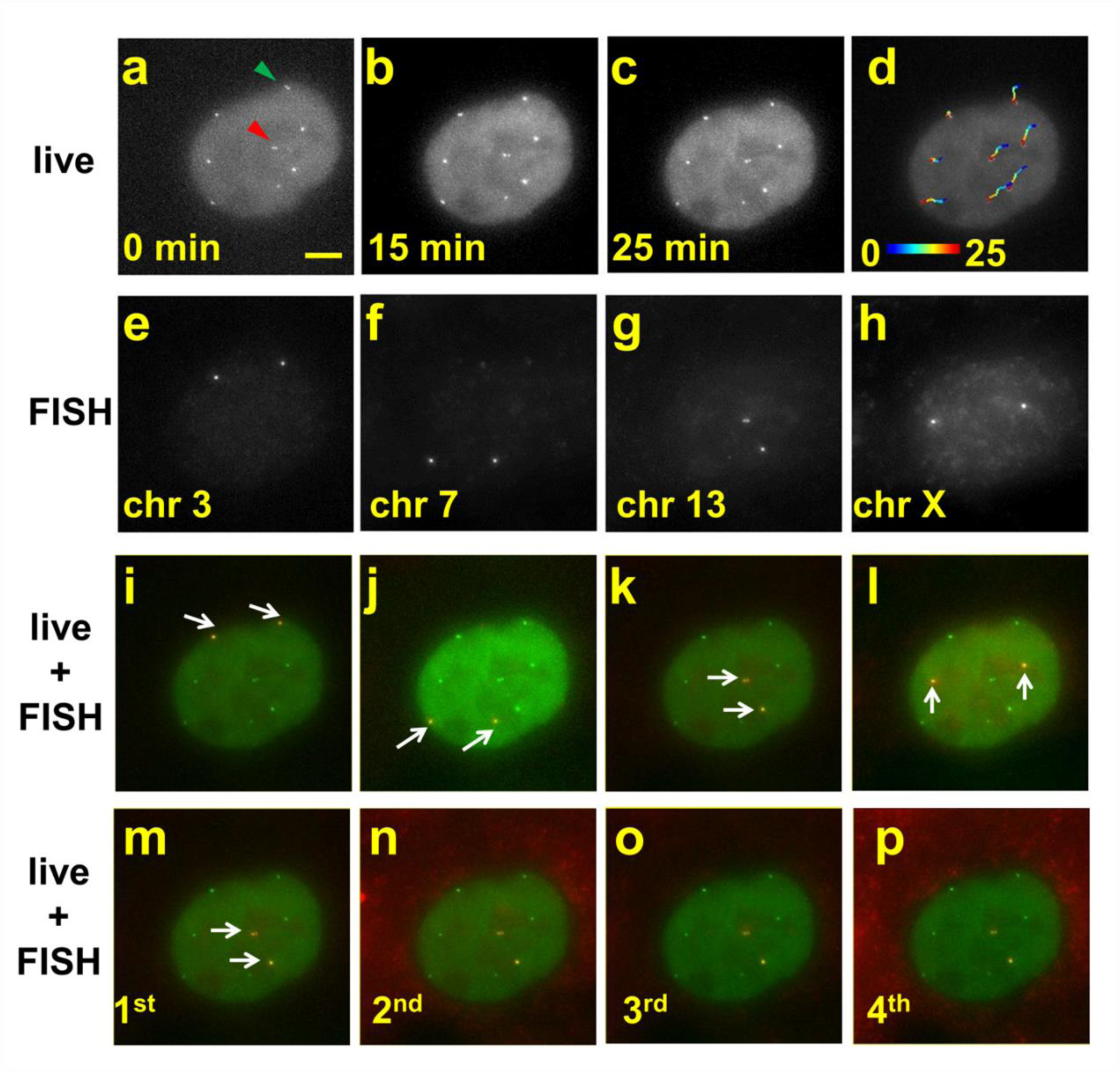
Sequential rounds of DNA FISH resolve identities of loci imaged in live-cell mode. (a-c) Representative micrographs of a cell from time-lapse images. 8 spots corresponding to 4 genomic regions in a diploid RPE cell are seen. Green arrowhead highlights a pair of sister chromatids that are distinguished at the beginning of the live imaging and came closer than the diffraction limit at later time. Red arrowhead highlights another pair of sister chromatids that are distinct throughout the image acquisition with fluctuating separation. (d) Overlay of the cell image at the end of the live observation with the corresponding loci trajectories. Color denotes time in minutes. (e-h) FISH images in four sequential FISH rounds reveal the identities of the loci. (i-l) Overlay of live images at 25 min with DNA FISH images from each round shows faithful registration and negligible nuclear deformation. The FISH spots in each round are highlighted with arrows. (m-p) Four sequential DNA FISH rounds staining Chr13 show consistent image registration between rounds. All micrographs here show the same image area and scale bar in (a) is 5 μm.

The cells are fixed at the end of the live observation and prepared for sequential DNA FISH. To correlate live imaging of genomic loci with sequential FISH, it is desirable to register the same area with sub-micrometer precision to correlate images between live condition and fixed FISH rounds. This task is challenging, especially since the denaturation of DNA duplex strands during DNA FISH preparation requires elevated temperatures; therefore, the sample has to be taken off the microscope stage. To address this issue, we use a combination of three measures to achieve faithful image registration. First, a 3D printed microscope stage adaptor ensures consistent sample orientation by restricting sample rotation (Fig. S1 in the Supporting Material). Second, we employ a rapid correlation-based algorithm to find the same cell observed in live imaging. Finally, a more sophisticated registration algorithm (17) that accounts for both translation and rotation is applied to register the live and fixed images based on the nucleus shape.

The identities of the four genomic loci are resolved in four rounds of DNA FISH as probes specific to each locus are sequentially introduced in each round (Figures 4e-h). Furthermore, overlay of the live-cell images and fixed-cell images demonstrate consistent loci position between live condition and fixed condition across various loci with a root mean square (RMS) error of ~52 nm in loci registration (Figure 4i-l; Fig. S11 in the Supporting Material). Similarly, we find images between sequential FISH rounds for a given probe register well with an RMS error of ~43 nm (Figure 4m-p; Fig. S12 in the Supporting Material). The successful image registration and principle component analysis on nuclear morphology (Fig. S13 in the Supporting Material) also confirms that negligible deformation is introduced during the FISH preparation steps. A closer inspection of Figure 4m-p shows that positions of sister chromatids and the distances between them are faithfully maintained in repeated sequential rounds and registered well with the live image, further confirming the method maintains almost intact nuclear morphology at both global and local scale and is compatible with high-resolution imaging to resolve fine chromatin structure.

### Multi-color live imaging enabled by correlative CRISPR imaging and sequential DNA FISH

Because the correlative CRISPR imaging and sequential DNA FISH uses a single color channel in live imaging to track multiple genomic regions, it opens up other color imaging channels in live cell imaging to extract information otherwise difficult to obtain. Here we demonstrated this capability by performing CRISPR imaging of 4 loci in cells expressing the Fucci probe, a widely used cell-cycle tracker which uses two colors to mark G1 or S/G2/M cell phase respectively (24). With the help from Fucci probe, we were able to distinguish G1 phase cells and early S phase cells (Figure 5a-c), both of which display singlet spots for each locus in CRISPR imaging results, while late S/G2 phase cells could identified by doublet spots corresponding to replicated sister chromatids (Figure 5d). Because the Fucci reporter itself already occupies two of the three color channels (green, orange and far red) that multi-color live cell imaging can be easily performed, this experiment is challenging for other multi-color CRISPR imaging methods. Potentially, with Fucci reporting the onset of S phase and CRISPR image detecting the replication of given genomic loci (e.g. based on intensity change of the labeled spot), we can profile replication timing and test how it affects the spatial organization of genome and the formation of topologically associated domains (TADs) formation (as recently proposed based on Hi-C results (25)). In addition, with other color channels opened up, our approach can also to be used to monitor the interaction of genomic loci with other nuclear components such as lamin, nuclear pore complex, and various nuclear bodies (26), many of which are known to be active genome organizers.

**Figure 5.**
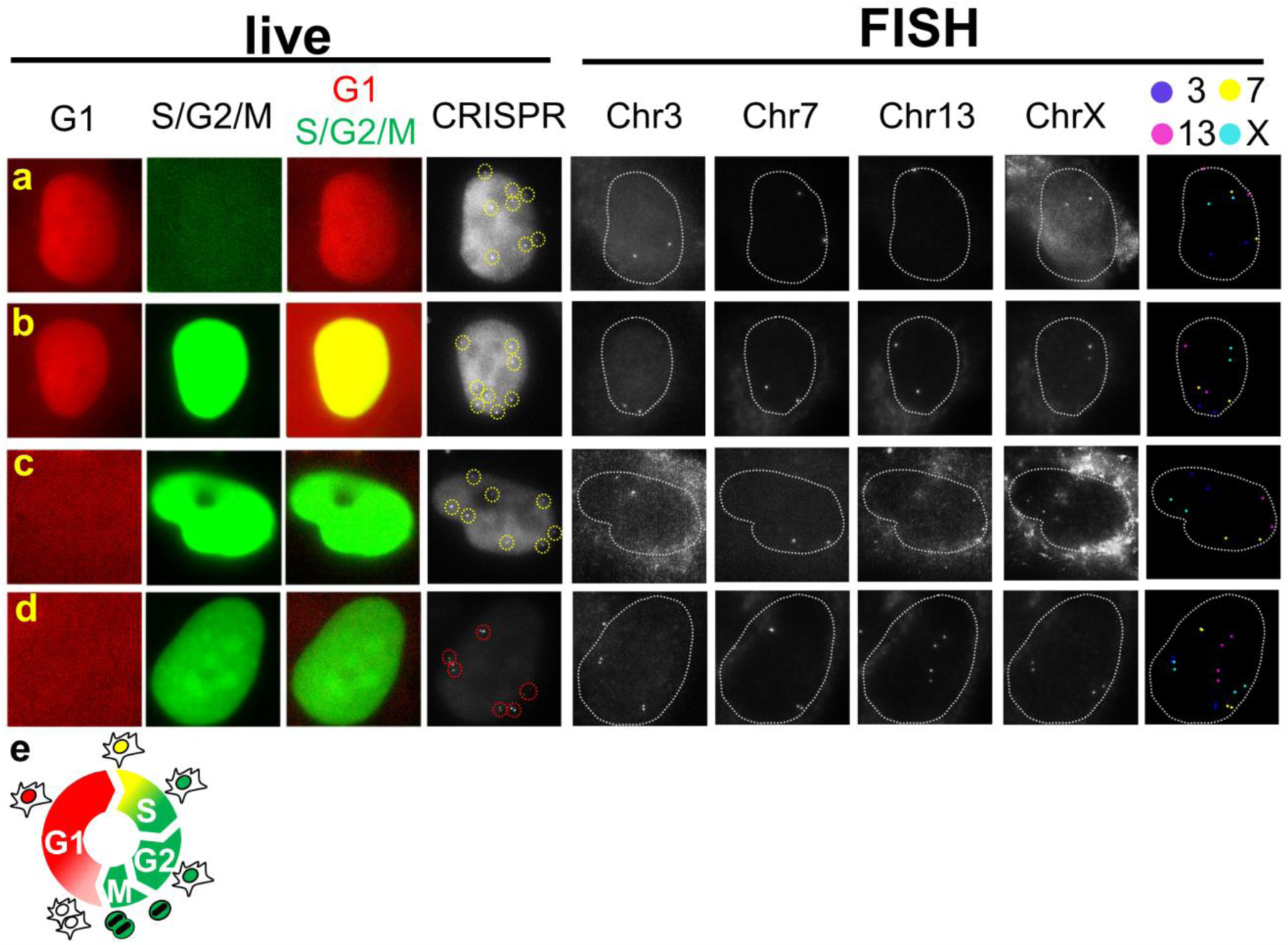
Simultaneous live CRISPR imaging of multiple genomic elements and cell cycle tracking. (a-d) Correlative imaging of CRISPR imaging and sequential DNA-FISH is performed on cells with Fucci cell cycle tracker at G1, onset of S, early S, and G2 phase respectively. The first three columns in live panel correspond to signals from Fucci cell tracker and the forth column in live panel shows CRISPR imaging where puncta are highlighted in circle. The first four columns in FISH panel are FISH images corresponding to four different genomic loci and the last columns in FISH panel shows all FISH puncta identified in sequential rounds in false-color. Nuclei are outlined with dotted line. (e) Schematic of Fucci cell cycle tracker.

## DISCUSSION

The current method thus combines the advantage of acquiring dynamics in live-cell imaging and multiplex imaging capacity in sequential FISH. It frees up other *in vivo* color channels for imaging applications such as RNA expression and processing (4, 27), protein expression (28, 29) and various nuclear components which are closely related to the spatial organization and dynamics of the genome. Moreover, a simpler system of live imaging with more uniform expression and assembly of Cas9 protein and sgRNA could potentially reduce system variability and facilitate further quantitative analysis. As DNA FISH has demonstrated the power to probe genome organization in high-throughput fashion such as HIP-map (30) and with super-resolution imaging using Oligopaints (31, 32), the current work could potentially add a new dimension of dynamic information. As the method of image registration between live and fixed conditions is directly transferrable to other systems, the concept of multiplex imaging through correlative imaging between live and fixed cells could be similarly applied to RNA imaging resolved by sequential RNA-FISH and protein imaging by sequential antibody staining.

## CONCLUSION

In summary, we introduce a correlative imaging method that combines the dynamic tracking capability of CRISPR imaging and the multiplicity of sequential FISH. After live imaging to obtain dynamics information of multiple genomic loci using one-color CRISPR system of one Cas9 protein and multiple sgRNAs, we perform rapid sequential rounds of DNA FISH to resolve loci identities. We also demonstrate a greatly simplified DNA FISH protocol that effectively stains genomic DNA in as short as 30 s in contrast to the common practice of overnight incubation. Our correlation-based algorithm to faithfully register between live images and fixed images can be readily adapted for other multiplex imaging applications.

## SUPPORTING MATERIAL

Thirteen figures, two text files, and one video are available.

## AUTHOR CONTRIBUTIONS

J.G. and B.H. designed research; J.G., H.L., and S.F. performed research; J.G., X.S, and H.L. analyzed data; and J.G. and B.H. wrote the article.

## ACKNOWLEDGMENTS

This work is supported by a W.M. Keck Foundation Medical Research Grant and the National Institute of Health (R21EB021453 and the Single Cell Analysis Program R33EB019784). H.L. receives the National Science Foundation Graduate Research Fellowship.

